# Anticipating emerging arboviruses: What comes after Zika?

**DOI:** 10.1101/143842

**Authors:** Michelle V. Evans, Courtney C. Murdock, John M. Drake

## Abstract

New vector-borne diseases have emerged on multiple occasions over the last several decades, raising fears that they may become established within the United States. Here, we provide a watchlist of flaviviruses with high potential to emerge in the US, identified using new statistical techniques for mining the associations in partially observed data, to allow the public health community to better target surveillance.

---

The start of the 21st century was marked by multiple emerging mosquito-borne diseases. West Nile virus was introduced to the eastern United States in 1999 and has since spread across the continental US, infecting tens of thousands of people and causing widespread mortality in bird populations (Hayes et al. 2005). A mutation in a strain of chikungunya virus increased that pathogen’s infectiousness in the invasive mosquito vector, *Aedes albopictus* (Asian tiger mosquito), leading to an expansion of the virus’s range from eastern Africa to Asia, and increasing the risk of disease transmission in temperate zones (Pialoux et al. 2007, Weaver and Forrester 2015). More recently, Zika virus has spread from Oceania to South America, and continued north at a rate exceeding 15,000 km/year (Zinszer et al. 2017), causing severe neurological diseases including infant microcephaly and Guillain-Barre syndrome(Lessler et al. 2016). In each case, the science and public health communities lacked basic knowledge about these previously geographically-limited diseases, and interventions were put in place only after the disease had become established. Our question is not *if* another mosquito-borne disease will emerge in the United States, but which will be next?

The sheer number of potential emerging pathogens underscores the infeasibility of comprehensive surveillance. Of over 530 known arboviruses that infect humans or domestic animals (Centers for Disease Control and Prevention 2016), most are poorly studied and knowledge gaps include information as basic as the species of mosquitoes that transmit the pathogen. Without a complete list of possible vectors, health agencies are unable to adequately plan for outbreaks or target virus surveillance. It is also difficult to predict whether an imported case of disease will initiate local transmission without knowing the efficiency at which mosquito species in the area can transmit the virus. Unfortunately, the development of diagnostic tools and vector competence studies (which are very labor intensive (Hardy et al. 1983) are often undertaken only after disease is endemic. But, without this information, health agencies are ill-equipped to prevent outbreaks.

To narrow the list of potential emerging diseases, we created a ‘watch list’ of flaviviruses (the family that includes well known arboviruses such as West Nile, dengue, yellow fever and Zika viruses) that might emerge in the United States. Our list is based on known associations of flaviviruses with mosquito species currently found in the US. Drawing on traits of known vector-virus pairs, in recent work, we developed a statistical model of trait combinations associated with the propensity of mosquito and virus species to form linked pairs (Evans et al. 2017). Known links were compiled from the literature on experiments showing which mosquito species can transmit the virus from an infected to a susceptible host. Fifteen traits of mosquitoes and twelve traits of viruses (e.g. geographic range, larval habitat, disease severity) were used to construct a predictive model. We then applied the resulting model to data on all the flaviviruses known to infect humans and the mosquito species found in the US.

The resulting network of potential virus-vector pairs in the United States (Fig. 1) contains thirty-seven mosquito species and twenty viruses, only five of which are currently or historically found in the US. Two viruses, Japanese encephalitis virus and Wesselsbron virus, have many more predicted links than the other viruses (36 and 29, respectively). Closely related to West Nile Virus, Japanese encephalitis virus infects over 67,000 people annually, with the majority of cases in Asia (Campbell et al. 2011). While only 1% of cases develop Japanese encephalitis, there is a 20 - 30% mortality rate among cases that progress to that stage (Kaur and Vrati 2003). Japanese encephalitis virus requires a non-human host to amplify the virus, domestic pigs, in its native Asia, but there are bird species in North America that are competent reservoirs for the virus (Nemeth et al. 2012). Much less is known about Wesselsbron virus, which has been historically restricted to a small geographic region of sub-Saharan Africa where it most commonly infects sheep and other livestock (Van Regenmortel et al. 2000). While symptoms of Wesselsbron virus in humans are relatively mild, it causes abortion in goats (Mushi et al. 1998) and cerebellar hypoplasia (a form of microcephaly) in calves (Coetzer et al. 1979) - a significant concern for the livestock industry.

**Figure 1:**
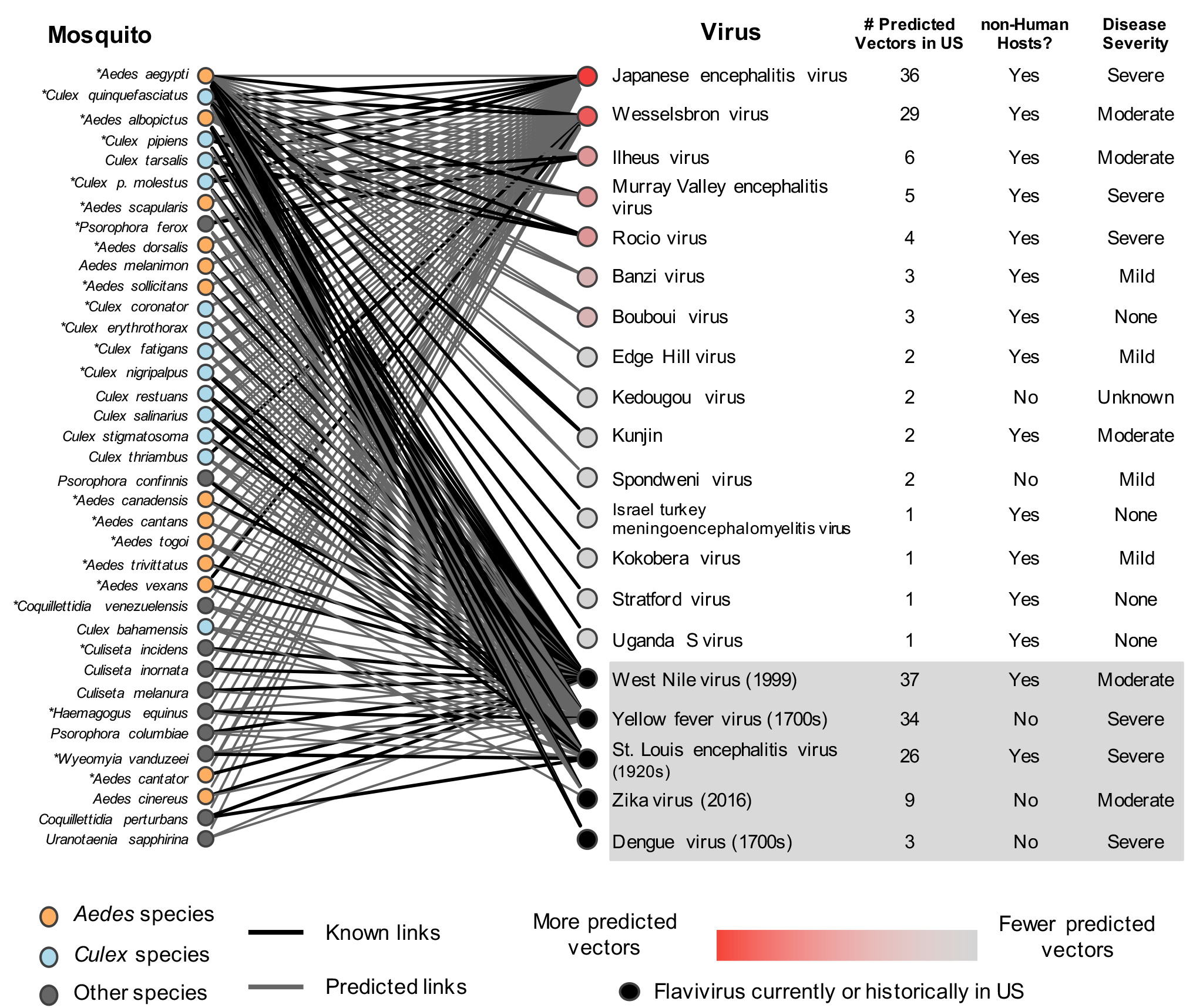
Network of flaviviruses known (black) or predicted (grey) to be transmitted by mosquito species found in the United States. Starred mosquitoes readily bite humans. Mosquitoes and viruses are rank ordered from top to bottom by the total number of possible links. Viruses that are currently or historically in the United States are shown in the gray box, with their approximate year of introduction, or first known case, in parentheses.

The virus with the most predicted links is West Nile virus (37 links). Other US-endemic viruses have between three and 34 predicted mosquito vectors. Yellow fever virus, which caused an outbreak in New Orleans as recently as 1905 (Crosby 2007), has a predicted 34 mosquito vectors. Since the introduction of the yellow fever vaccine in the mid-20th century, transmission has been mostly restricted to the African continent (Frierson 2010), however, the 2016 epidemic in Angola demonstrates the inadequacy of current immunization regulations (Barrett 2016). By contrast, dengue virus, which is locally transmitted in Puerto Rico and the Gulf Coast, is only predicted to be vectored by three mosquito species endemic to the United States. Despite the small number of competent vector species, there were nonetheless an estimated 100,000 dengue cases in the continental US in recent years (Hotez 2008). The high number of cases highlights the importance of the identity of predicted vectors on disease transmission, in this case, *Aedes aegypti,* a mosquito species that is well-adapted to live with humans and is a highly efficient vector of many human pathogens (Powell and Tabachnick 2013).

These viruses illustrate, but do not exhaust, the risk of emergence posed by viruses circulating elsewhere in the world. With the rise in globalization and availability of air transportation, a mosquito-borne pathogen can travel across the world in less than 24 hours, connecting countries with active transmission to countries without disease (e.g. Zika and chikungunya (Escobar et al.2016). While several states have large mosquito surveillance programs, most do not (Hamer 2016), and the accompanying virus surveillance programs screen for a limited number of pathogens. The emergence of Zika virus in the United States, and the corresponding scramble to control its spread (Halford 2016, Tinsley 2016), highlights the need to shift from reactive to proactive surveillance and control strategies for mosquito-borne diseases (Olival and Willoughby 2017). Focusing on the viruses with a high potential to emerge could improve the efficiency and reliability of surveillance without overly burdening public health institutions and help guide future vaccine development, such as the Coalition for Epidemic Preparedness Innovations (Butler 2017). Finally, the approach we present here can be applied to other disease systems to identify groups of viruses and bacterial pathogens most likely to emerge in the United States or any other country.

